# Are ant-plant mutualisms stronger at lower latitudes? A case study using the castor plant in the Indian subcontinent

**DOI:** 10.1101/2025.01.07.631652

**Authors:** Pooja Nathan, Arshyaan Shahid, Anoushka Datta, Vinita Gowda, Megan E. Frederickson

## Abstract

The Biotic Interactions Hypothesis (BIH) predicts that species interactions will intensify from the poles to the tropics. We studied the ant-plant defensive interaction mediated by extrafloral nectaries (EFNs) in the castor bean plant, *Ricinus communis,* across a ∼21° latitudinal gradient in its naturalized range in India. Among our study sites, we found that the ant-EFN mutualism in castor bean gets stronger from sub-tropical to tropical latitudes. Investment in mutualism by plants and visitation by ants increased from higher to lower latitudes. Further, latitude significantly explained ant community composition. However, contrary to the predictions of the BIH, standing herbivory increased with latitude, perhaps because plants invest less in biotic defence at higher latitudes. To our knowledge, our study is the first to test for the patterns predicted by the BIH for an ant-plant interaction in the Paleotropics. Results from our study can also help inform sustainable pest control practices for *R. communis* since India is the world’s largest producer of castor oil.

## Introduction

The tropics harbour most of the world’s biodiversity (Gaston, 2000) and therefore most of the world’s species interactions. In his well-known essay “Evolution in the tropics,” Dobzhansky (1950) famously proposed that the main challenge that temperate organisms face is an unfavourable climate, while “the challenges of tropical environments stem chiefly from the intricate mutual relationships among inhabitants.” This idea has become known as the Biotic Interactions Hypothesis (BIH), which posits that the primary agents of selection vary from temperate regions to the tropics: selection is primarily driven by abiotic factors in temperate regions, but biotic factors, i.e., interactions with other species, play a stronger role in determining fitness in the tropics (Dobzhansky, 1950; Schemske et al., 2009). Therefore, the BIH predicts that species interactions will be more intense or specialized towards the tropics (Schemske et al., 2009). The BIH figures prominently in some (but not all) explanations for why biodiversity increases from the poles to the tropics (Brodie & Mannion, 2023; Schemske et al., 2009; Willig et al., 2003), for example in addressing whether biotic interactions contribute meaningfully to more rapid speciation at lower latitudes by becoming more specialized or stronger from the poles to the tropics (used interchangeably with “lower latitudes” in this study). Addressing this question is critical for evaluating evolutionary mechanisms that might contribute to the latitudinal gradient in species richness.

Empirical support for the BIH has been mixed. There is evidence both for and against higher specialization in the tropics (Forister et al., 2015; Schleuning et al., 2012; Vázquez & Stevens, 2015). However, studies across study systems and interaction types suggest that the effect size of interspecific interactions is stronger for both species at tropical latitudes in comparison to temperate latitudes (Baskett et al., 2020; Baskett & Schemske, 2018; Hargreaves et al., 2019; Roslin et al., 2017 but see Freeman et al., 2022; Lim et al., 2015). This pattern is attributed to the higher abundance of interacting partners in the tropics leading to increased encounter rates, a higher diversity of interacting species due to high species richness at low latitudes, or both, regardless of the type of interaction. Alternatively, less seasonal variation in climate at lower latitudes may enable year-round interdependencies of interacting species leading to stronger interactions (Schemske & Mittelbach, 2017).

Most studies testing the BIH have focussed on antagonistic interactions such as herbivory, granivory, or predation (Freeman et al., 2019; Hargreaves et al., 2019; Moles et al., 2011; Roslin et al., 2017; Zvereva & Kozlov, 2021). However, the BIH may also be applied to mutualistic interactions, for example in the prediction that an organism at lower latitudes with stable abiotic conditions may be more likely to encounter a compatible mutualistic partner since they are more abundant or diverse. Thus, per the BIH, positive interactions are likely to be more beneficial, and negative interactions more detrimental as we move from temperate to tropical latitudes. Studies investigating patterns in mutualistic interaction strength across latitudes are much more scarce than those for antagonisms (Baskett et al., 2020; Gorostiague et al., 2023; Harrison et al., 2024). To bridge this gap, we studied the variation of ant-plant defensive mutualism across latitude in the Indian Subcontinent.

Ant-plants mutualisms serve diverse functions and span the generalization-specialization continuum in ecosystems across the world. In particular, tropical biomes are home to many iconic plant-ant mutualisms, such as the ant acacias in African savannas (Stanton et al., 1999) and Central America (Janzen, 1966), plant-farming ants in Fiji (Chomicki & Renner, 2016), and “devil’s gardens” in the Amazonian rainforest (Frederickson et al., 2005). Ant-plant mutualisms mediated by extrafloral nectaries (EFNs) or domatia (structures produced by plants to house predatory arthropods like ants or mites) are more prevalent in the tropics than in temperate regions, and this is likely driven by high ant species richness and environmental variables such as climate. Yet, few studies have investigated if ant-plant mutualisms show a latitudinal gradient in their specialization or strength. Both extrafloral nectaries and domatia constitute an indirect defence mechanism in plants, in which the plants offer a nectar reward (in EFNs) or nesting space (in domatia) to potentially protective ants (Bronstein et al., 2006; Heil & McKey, 2003). Ant-domatia interactions are known to be quite specialized in comparison to ant-EFN interactions which are highly generalized (Chomicki & Renner, 2015; Heil, 2008). Thus, compared to domatia, EFNs may experience fewer constraints and may be more likely to vary with the environmental variables that covary across latitudes.

We studied an ant-plant mutualism across a wide (∼21 degree) latitudinal range in the Indian subcontinent to test if the strength of this interaction increases with decreasing latitude. We did this by quantifying host plant investment in mutualism and ant community characteristics in plant populations spanning a latitudinal gradient, while also measuring standing herbivory rates. To do this, we studied the mutualism between *Ricinus communis* (Euphorbiaceae), the castor bean plant, and the ant species visiting its EFNs in its naturalized range in India. In addition to being agriculturally important as an oilseed crop (Anjani, 2014, p. 201; Patel et al., 2016), *R. communis* is notable for its interactions with a range of herbivores and mutualists (López-Guillén et al., 2020; Waters et al., 2014), and for its long-standing association with humans. Having originated in eastern Africa, *R. communis* was domesticated thousands of years ago by humans who extracted oil from its seeds. It has been proposed that the use of castor oil by humans very likely contributed to its arrival in much of its modern naturalized and introduced ranges (Polito et al., 2019; Xu et al., 2021). The seeds of this species also contain the potent toxin ricin, a lectin, likely a generalized anti-herbivore defence chemical (Audi et al., 2005; Olsnes, 2004). Castor bean plants produce EFNs on the petioles of leaves and near inflorescences (Figs. 1c, 1d) that attract ants and other primarily hymenopteran visitors (Nathan et al., 2023; Waters et al., 2014). The presence of ants likely wards off potential herbivores since they may bite, sting or spray acid. Results from a common garden experiment (Nathan et al., in prep) showed that *R. communis* individuals subjected to ant exclusion treatments suffered increased herbivory (Fig. S1), indicating a resource-protection mutualism between the ants and plants.

**Figure 1:**
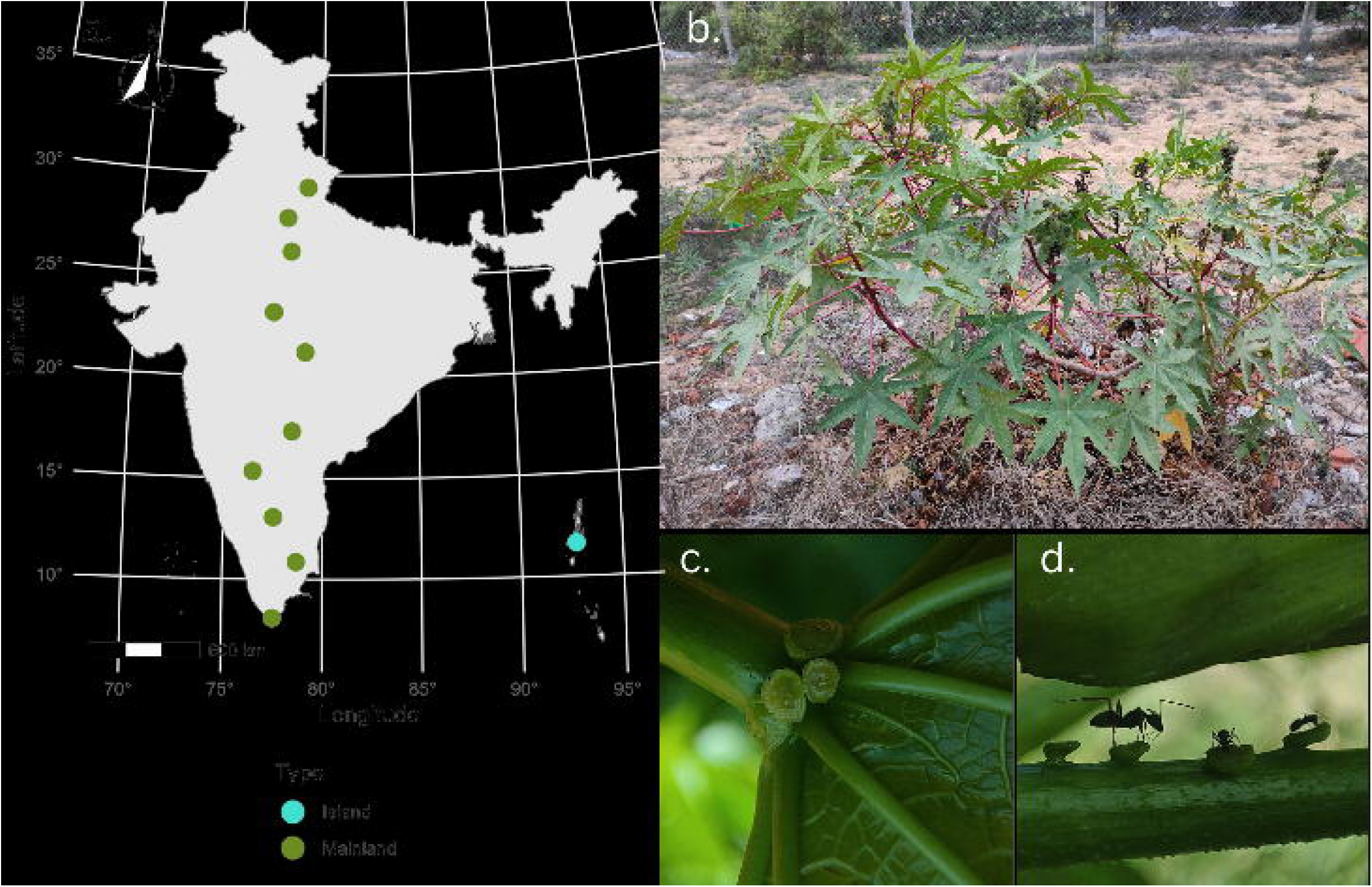
Study site and study system. a) Map of sites sampled across the Indian subcontinent. Mainland sites are coloured green and the island site is coloured blue. b) *Ricinus communis*, the castor oil plant, in the southernmost site, Kanyakumari. c) A *R. communis* leaf base EFN d) *Ricinus communis* petiole EFNs being visited by ants. Photos: Pooja Nathan.

To assess allocation to EFNs by plants and visitations to the EFNs by ants, we measured plant traits in *R. communis* populations and characterised the ant visitors to EFNs across a latitudinal gradient. We asked: a) Do plants invest more in EFNs towards lower latitudes? b) Does the abundance and diversity of ant partners visiting *R. communis* EFNs increase from higher to lower latitudes? c) Does latitude predict herbivory on *R. communis*? Our study presents a novel test of the BIH for ant-plant interactions and herbivory in a region of the world that is underrepresented in the ecological and biogeographic literature.

## Materials and methods

### Study system and study sites

We sampled mutualism-related traits, interactions with ants, and herbivory in eleven populations of *R. communis* across the Indian subcontinent between April and June 2022 (Fig. 1a). *Ricinus communis* is ruderal and closely associated with human settlements, therefore all our sample sites were close to urban areas. To cover the latitudinal extent of India without varying the longitude across sites, we identified ten Indian cities that fall around a longitude connecting the northernmost and southernmost points of the country (Fig. 1a). Our northernmost site was Ramnagar (in the state of Uttar Pradesh; 29.3948° N, 79.1265° E) and our southernmost site was Kanyakumari (in the state of Tamilnadu, 8.0844° N; 77.5495° E), spanning a latitudinal range of around 21°. Further, we sampled *R. communis* in Port Blair (11.6234° N, 92.7265°E; Fig. 1a), the capital city of the Andaman and Nicobar Islands, an archipelago located in the Bay of Bengal on the east coast of the Indian peninsula, since interaction dynamics often vary in islands in comparison to the mainland (Delavaux et al., 2024; Freedman et al., 2024). To control for the interaction between latitude and seasonality in climate, we randomized the order in which the sites were sampled (i.e., we did not sample sites from north to south, or from south to north). At each site, we identified a population of *R. communis* plants surrounded by vegetation, away from the urban core where ant diversity is likely low (Buczkowski & Richmond, 2012, but see Perez & Diamond, 2019). *Ricinus communis* is considered a weedy species and is often removed periodically, thus we identified and tagged 10-20 plants in each site that had no signs of human damage (see Table S1 for the number of plants sampled at each site). We chose plants that were at least 1 m away from each other to sample unique genets. Since it is a perennial, *R. communis* can take on a tree-like habit, thus, to control for the effects of ontogeny, all plants sampled were under 2m in height.

### Plant traits

We used EFN counts and areas to measure investment in mutualism by plants across sites. *Ricinus communis* produces many distinct types of EFNs, and we considered two unique types in this study. Leaf base EFNs (Fig. 1c) are found at the point of attachment of the stem to the petiole, and petiole EFNs (Fig. 1d) are found all over the petiole and stem (Fig. 1c, 1d). The youngest parts of the plant tend to be highly defended as per the predictions of Optimal Defence Theory (Calixto et al., 2021; McKey, 1974), thus on each plant, we tagged a young branch with 3-4 leaves and an apical bud and counted the number of leaf base (LB) and stem and petiole (S) EFNs. These counts were standardised by the number of leaves on the branch to obtain the number of EFNs of each type per leaf. We measured areas of leaf base EFNs and stem EFNs from photographs that included a ruler for calibration using ImageJ (Schneider et al., 2012). To measure nectar volumes, we restricted access of ants to the experimental young leaf for 24 hours by applying a layer of tape to the petiole, over which we painted a thick layer of Tree Tanglefoot. We then photographed the drops from above and the side to estimate the volume of each drop through image analysis using the scale in the images. We were able to obtain only a small number of direct volume measurements by collecting extrafloral nectar using capillaries due to the extremely small size of EFN droplets and instances of nectar robbery by airborne insects like flies. Nonetheless, EFN volume was strongly correlated with nectary area for both EFN types (Fig. S2), thus we used nectary areas as a correlate of investment in mutualism. To obtain a snapshot of plant fecundity, we counted the number of fruits on each plant we sampled. Finally, we measured the width of the stem at 10cm above the ground for each plant as a proxy for plant age.

### Species interactions

At a total of 11 sites, we scanned and quantified all tagged plants between 8 am and 12 pm on 2 to 4 consecutive days to characterize a) the abundance and diversity of ant visitors on stem and leaf-base EFNs and b) the levels of standing herbivory. On each experiment day, we scanned all plants at least five times. We sampled all ant visitors on the three topmost fully open leaves of a young branch. Most ant species, with one exception, were identified in at least to genus (Table S2). Ant abundance for each species was standardized by the number of leaves to obtain the number of ants of each species per leaf. To quantify standing herbivory, we selected 10 leaves at random on each plant and calculated the proportion of leaves showing visible evidence of herbivory (i.e., folivory, leaf miner tracks, etc.). If plants had fewer than 10 leaves, we scanned all leaves for signs of herbivory and calculated the proportion accordingly.

### Statistical analysis

All statistical analyses were carried out using R version 4.4.1 (R Core Team). We used multivariate linear models (*mlm* function) to understand the effects of latitude on EFN counts and areas, with stem width and site type (island vs. mainland) as covariates. Using the *lm* function (R Core Team), we ran one model each for EFN counts and EFN areas, jointly specifying the counts or areas of both leaf base and petiole EFNs as response variables. We used linear models using the *lm* function to understand the effect of latitude on herbivory and ant abundances. For herbivory, we included stem width, ant abundance, and site type as covariates.

We used the *vegan* package (Oksanen et al., 2024) to understand how ant species richness, diversity, and community composition vary across latitudes. Since the cumulative number of ant species largely stabilized by the 6^th^ scan (Fig. S3a) but increased with the number of scan days (Fig. S3b), we calculated the mean abundance across the first six scans on each sampling day, for all experiment days for which data were available. To investigate the relationship between latitude and species richness, we computed ant species richness for each site using the *specnumber* function and we used a linear model (*lm*) to estimate the effect of latitude on species richness across sites. Similarly, we used the *diversity* function to compute the Shannon diversity of ant species for each site and regressed these values against latitude using a linear model. We used Non-metric Dimensional Scaling (NMDS) to visualise how ant community composition changes across sites using the *metaMDS* function implemented on Bray-Curtis dissimilarities between ant species abundances across sites. Finally, to investigate if the effect of latitude on ant species composition is statistically significant, we ran a permutational multivariate analysis of variance (PERMANOVA) with Bray-Curtis distances using the *adonis2* function.

We used a Structural Equation Model (SEM) implemented with the *lavaan* package (Rosseel, 2012) to further investigate the relationships between different variables in our dataset. We specified an a priori model outlining predicted relationships between variables representing mutualism traits (EFN counts of leaf base and petiole EFNs), plant age, herbivory, ant abundance and the number of fruits as a snapshot measure of fitness, as shown in Fig. 4a. In addition to including a direct effect of EFN on fitness, we also included an indirect effect of EFNs where EFNs may affect plant fitness by increasing ant abundance, which may decrease herbivory.

Finally, we included a covariance term between leaf base and petiole EFNs. We excluded EFN areas from this analysis since we had fewer observations of this variable and including them in the model significantly reduced statistical power. Since our variables spanned orders of magnitude, we scaled and centered them using the *scale* function before running the SEM using the *sem* function.

## Results

### Plant traits

We found a latitudinal gradient in mutualistic traits and interactions. As predicted, the number of leaf-base EFNs significantly decreased with increasing latitude (Fig. 2a, Table 1A). However, this trend was not observed for petiole EFNs, which showed weak positive but non-significant variation with increasing latitude (Fig. 2b, Table 1A). Stem width, used as a correlate of plant age, had a marginally significant negative effect on the number of both types of EFNs (Table 1A), meaning plants may have more EFNs when they are younger. The areas of both types of EFNs decreased significantly with increasing latitude (Figs. 2c, 2d, Table 1B), with southern sites having larger EFNs. We observed a strong positive effect of latitude on the proportion of leaves showing evidence of herbivory (Fig. 2e, Table 1C) and northern sites experienced higher herbivory, contrary to our prediction. Furthermore, we found that ant visits decreased significantly with increasing latitude (Fig. 2f, Table 1C), with plants from more southern sites attracting more ants. The island site did not differ significantly from the mainland sites in terms of EFN counts, areas, or ant visits, but mainland plants suffered more herbivory than the island plants (Figs. 2a-2f insets, Table 1C). Taken together, these results suggest that in *R. communis,* mutualism, viewed from both the plant and ant perspectives, intensifies as we move from temperate to tropical latitudes. On the other hand, herbivory weakens.

**Figure 2:**
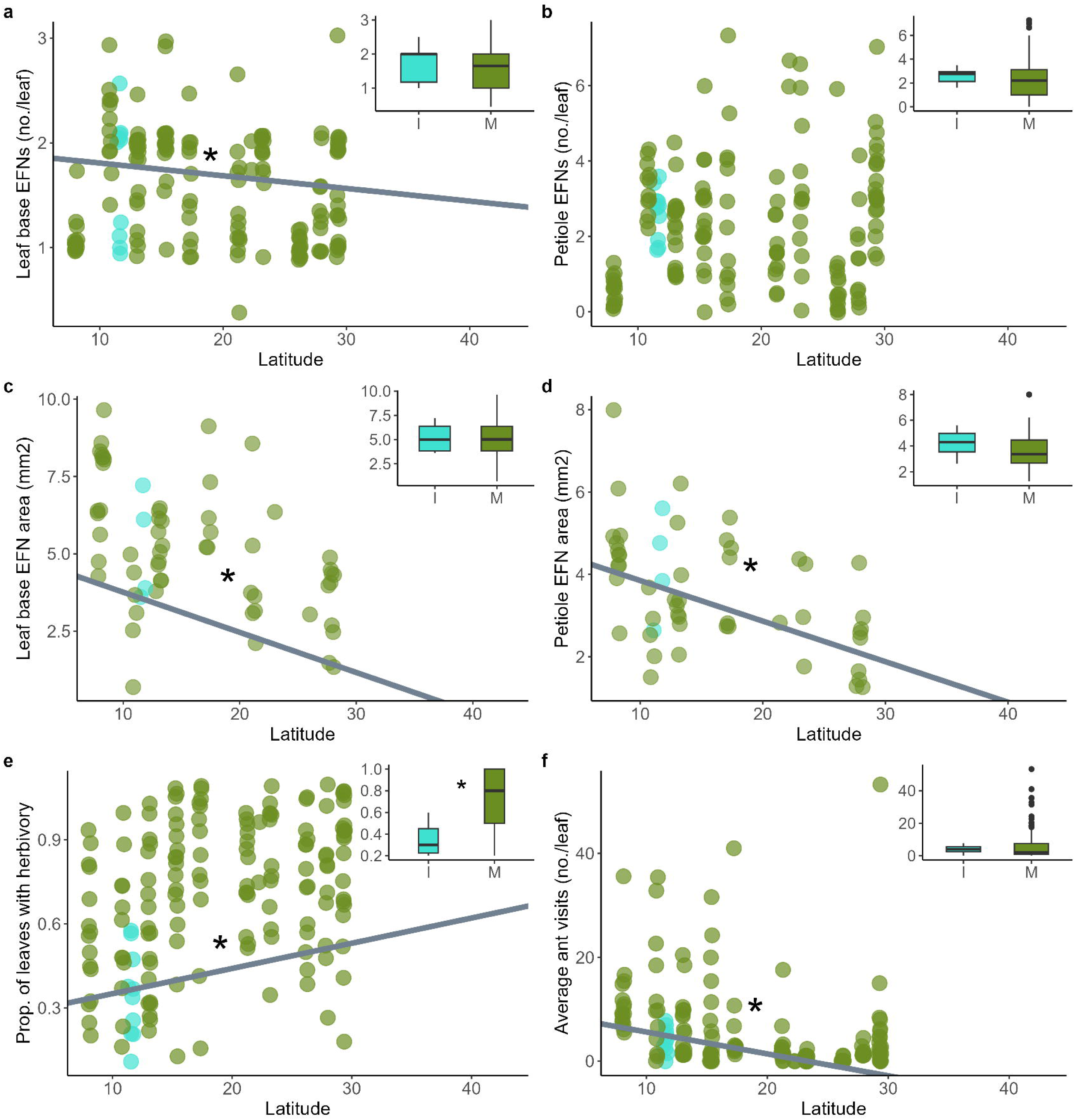
Variation in mutualism traits, herbivory, and ant visits across latitude. Blue points represent the island site (Port Blair) and green points represent the 10 mainland sites. The inset plots compare the variables shown in the plots between mainland and island sites, and I = Island, M = Mainland. Significant relationships are indicated by an asterisk. a) Variation in the number of leaf base EFNs across latitude. c) Variation in the area of petiole EFNs across latitude. d) Variation in the area of leaf base EFNs across latitude. e) Variation in the proportion of leaves showing herbivory across latitude. f) Variation in the average number of ants per leaf across latitude. See Table 1 for the complete results of statistical models.

**Table 1:** Results of linear models. A) Multivariate linear model results for the effect of latitude, plant age, and mainland vs. island location on leaf base and petiole EFN counts. B) Multivariate linear model results for the effect of latitude, plant age, and mainland vs. island location on leaf base and petiole EFN areas. C) Linear model results for herbivory and ant visits. Significant effects are italicized and indicated by *** for p ≤ 0:001, ** for p ≤ 0:01 and * for p ≤ 0:05. Marginally significant effects are indicated by a period (.).

| **A. Multivariate linear model - EFN counts** |  |  |  |  |
| --- | --- | --- | --- | --- |
| Leaf base EFN counts |  |  |  |  |
| **Coefficients** | **Estimates** | **SE** | **t value** | **p-value** |
| *Intercept* | 1.92904 | 0.26341 | 7.323 | 1.29e-11 *** |
| *Latitude* | -0.01213 | 0.00601 | -2.017 | 0.0455 * |
| **Plant age** | -0.01919 | 0.011211 | -1.712 | 0.0890 . |
| **Mainland vs. Island** | 0.071918 | 0.242697 | 0.296 | 0.7674 |
| Petiole EFN counts |  |  |  |  |
|  | **Estimates** | **SE** | **t value** | **p-value** |
| *Intercept* | 2.95035 | 0.81146 | 3.636 | 0.000378 *** |
| **Latitude** | 0.02717 | 0.01852 | 1.467 | 0.144449 |
| **Plant age** | -0.06075 | 0.03454 | -1.759 | 0.080557 . |
| **Mainland vs. Island** | -0.60338 | 0.74765 | -0.807 | 0.420896 |
| **B. Multivariate linear model - EFN areas** |  |  |  |  |
| Leaf base EFN areas |  |  |  |  |
|  | **Estimates** | **SE** | **t value** | **p-value** |
| *Intercept* | 5.05809 | 1.89242 | 2.673 | 0.01112 * |
| *Latitude* | -0.1298 | 0.03926 | -3.306 | 0.00211 ** |
| **Plant age** | 0.04621 | 0.06898 | 0.67 | 0.50711 |
| **Mainland vs. Island** | 1.67779 | 1.81944 | 0.922 | 0.36243 |
| Petiole EFN areas |  |  |  |  |
|  | **Estimates** | **SE** | **t value** | **p-value** |
| *Intercept* | 4.83484 | 1.39476 | 3.466 | 0.00135 ** |
| *Latitude* | -0.09848 | 0.02894 | -3.403 | 0.00161 ** |
| **Plant age** | 0.02034 | 0.05084 | 0.4 | 0.69133 |
| **Mainland vs. Island** | 0.0249 | 1.34097 | 0.019 | 0.98529 |
| **C. Linear models - herbivory and ant visits** |  |  |  |  |
| Herbivory |  |  |  |  |
|  | **Estimates** | **SE** | **t value** | **p-value** |
| *Intercept* | 0.26099 | 0.11989 | 2.177 | 0.031037 * |
| *Latitude* | 0.00899 | 0.00286 | 3.143 | 0.002010 ** |
| **Plant age** | -0.00649 | 0.005021 | -1.293 | 0.19793 |
| **Ant visits** | -0.0036 | 0.002345 | -1.536 | 0.126563 |
| *Mainland vs. Island* | 0.36892 | 0.10914 | 3.38 | 0.000921 *** |
| Ant visits |  |  |  |  |
|  | **Estimates** | **SE** | **t value** | **p-value** |
| *Intercept* | 9.81496 | 4.05615 | 2.42 | 0.0167 * |
| *Latitude* | -0.4213 | 0.09258 | -4.551 | 1.08e-05 *** |
| **Plant age** | -0.1528 | 0.17264 | -0.885 | 0.3775 |
| **Mainland vs. Island** | 5.36891 | 3.73719 | 1.437 | 0.1529 |

### Ant community analysis

Across sites, we found 21 ant species visiting the EFNs of *R. communis* (Table S2). Both ant species richness and Shannon diversity tended to decrease with increasing latitude, but the trends were not statistically significant (Fig. 3a, 3b, Table S3). Because processes driving ant community assembly on the island may differ from those on the mainland, we excluded the island site (Port Blair) from analyses but still found non-significant relationships between both species richness and Shannon diversity and latitude (Table S3). PERMANOVA scores calculated on Bray-Curtis dissimilarity distances between sites revealed that latitude significantly predicted ant communities (Pseudo-F = 2.0756, R^2^ = 0.187, p = 0.049) and explained around 19% of their variation (Fig 2c). When the island site was excluded from the model, latitude significantly and more strongly predicted ant communities (Pseudo-F = 2.3363, R^2^ = 0.226, p = 0.028) and explained around 23% of their variation. Taken together, these results show that the ant communities that visit *R. communis* change in composition, but not diversity, with latitude.

**Figure 3:**
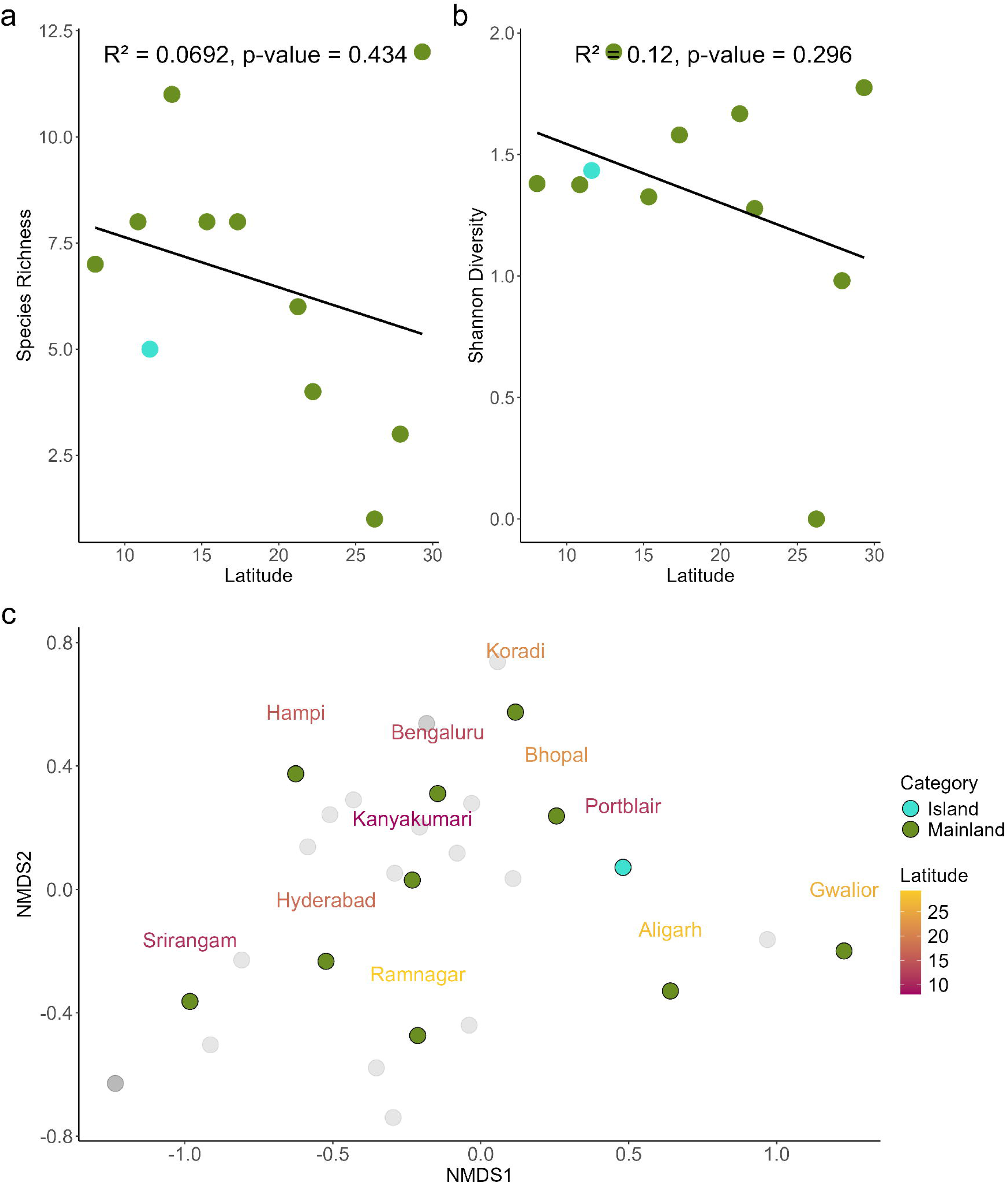
Ant community characterization of sites. Blue points represent the island site (Port Blair) and green points represent the 10 mainland sites. a) Variation in species richness across latitude. b) Variation in Shannon diversity with latitude. c) NMDS plot showing the ant species (grey dots) and sites (blue or green dots). The names of the sites are coloured by latitude, with darker red colours indicating lower latitudes. See Table S3 for the results of statistical models of species richness and Shannon diversity.

### Structural equation modelling

The global model showed a good fit with a Comparative Fit Index (CFI) of 1 and Tucker-Lewis Index (TLI) of 1.089. We found significant support for many paths outlined in the a priori model (Fig. 4a). As borne out by the linear models, latitude positively influenced herbivory, and negatively influenced ant visitation and the number of petiole EFNs (Fig. 4b, Table S4). Further, the number of fruits on plants, a snapshot of plant fitness when sampled, also declined with increasing latitude (Fig. 4b, Table S4). Stem width, used as a correlate of plant age, had a significant positive effect on the number of fruits, suggesting that older and more mature plants produced more fruits (Fig. 4b, Table S4). However, we found no evidence for direct or indirect causal links between EFN counts, herbivory or ant visitation and the snapshot count of the number of fruits. We found that leaf base EFNs, but not petiole EFNs, had a significant positive effect on ant visitation, suggesting that indirect defence by ants is mediated by EFNs (Fig. 4b, Table S4). Finally, we found a significant positive covariance between leaf base and petiole EFN counts, suggesting a common developmental mechanism.

**Figure 4:**
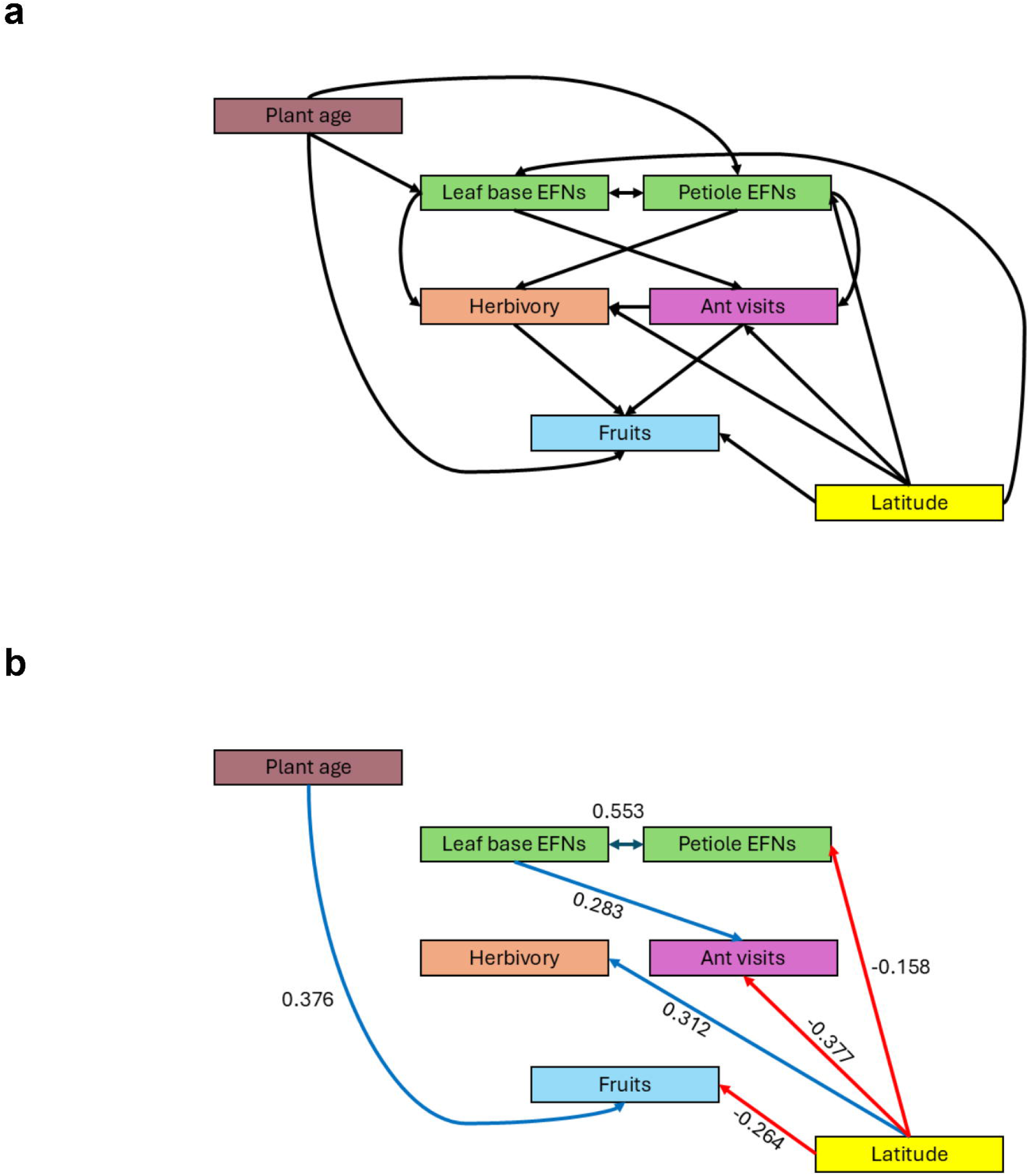
Structural equation models. a) A priori model showing all expected relationships. b) Statistically significant relationships. Positive relationships are shown with blue arrows, and negative relationships are shown by red arrows. Standardized estimates and covariances are shown alongside the arrows. For the complete results of the SEM, see Table S4.

## Discussion

As per the predictions of the BIH, the ant-plant mutualism in *Ricinus communis* intensifies from temperate to tropical latitudes in the Indian subcontinent. We found evidence for this in both partners: *R. communis* produced higher numbers and larger nectaries in more southern sites, and plants in more southern sites were visited by more ants than plants at higher latitude sites. Contrary to the predictions of the BIH, we observed that herbivory intensified towards the north, which aligns with a weakening of this defensive mutualism at higher latitudes. We found that latitude significantly predicted ant community composition, but not ant species richness or diversity, across sites. Finally, we found that EFNs increase ant visits, but snapshot counts of ant visits do not predict standing herbivory rates.

Studies focused on ant-EFN interactions have shown that it is generalized and may not require a deep coevolutionary history between the ant and plant partner for successful interaction to occur(Heil & McKey, 2003; Lange et al., 2013). Global analyses show that the diversity of ant species increases from high to low latitudes and it is likely that this trend is reflected in the sites sampled across decreasing latitudes within India (Dantas & Fonseca, 2023). Thus, *R. communis* individuals growing in southern latitudes may encounter more diverse ants, increasing the probability that some of them are mutualistic and protect the plants from herbivore attacks. In the presence of herbivory, individuals producing high volumes of EFN may be selected in southern sites, since they are more likely to attract effective ant bodyguards. Thus, in this case, the latitudinal pattern in mutualism may have been caused by the latitudinal diversity gradient in ant species richness. This may also explain the lack of a direct causal link between ant abundance and herbivory in the SEM. If only a subset of the ants attracted by EFNs contributed to effective indirect defence, the total number of ants visiting a plant may not be a good predictor of the defensive services obtained. Latitude significantly predicted ant community composition across sites, and southern sites may have more mutualistic ant communities which ultimately provide improved defensive function compared to northern sites. Further, since our measurements of herbivory and ant visitation were for only a single snapshot in time, and standing herbivory represents past damage while the number of ants on a plant represents only current visitation, it is also possible that total ant visitation does directly affect the rate of herbivory when measured over more comparable time windows.

We found that for both leaf base and petiolar EFNs, nectary areas decreased with increasing latitude. However, only leaf base EFN counts decreased with increasing latitude based on the results of the linear models. Further, based on the results of the SEM, leaf base EFN counts, but not petiole EFN counts influenced ant abundance. Finally, the results of the SEM suggest that the number of petiole EFNs decreases with increasing latitude, with latitude negatively affecting leaf base EFN counts, but only indirectly via petiole EFNs, since they are significantly positively correlated (Fig. 4b). Taken together, these results suggest that the leaf base and petiolar nectaries may serve different biological functions and are likely governed by differing processes, although they do have shared developmental pathways as evidenced by a significant covariance. Different types of extrafloral nectaries within the same plant species are known to vary in structure, induction, and nectar composition (Chatt et al., 2021; Escalante-Pérez et al., 2012). However, little is known about the ecological significance of this variation. Our SEM results suggest that in *R. communis*, leaf base EFNs may play a more direct role in attracting ants than petiole EFNs.

Leaf base EFNs are often larger than petiole EFNs in *R. communis*, and from a foraging perspective are likely to provide larger nectar rewards to ant visitors thus potentially attracting more ants. Further, it is known that EFN secretion can be induced by herbivore damage in *R. communis* (Wäckers et al., 2001) and since the majority of herbivores tend to eat leaves, at least when plants are primarily vegetative, leaf base EFNs may be more likely to get induced than petiole EFNs. More studies of the nectar production, composition and ant communities visiting different types of nectaries when faced with herbivore damage are needed to understand these mechanisms.

The predictions of the BIH involving plant systems have mostly been tested in latitudinal gradients of herbivory. According to the BIH, a higher abundance and diversity of herbivores in tropical latitudes will contribute to increased rates of herbivory. Studies that have attempted to quantify rates of herbivory across latitude have found mixed results; some studies show an increase in herbivory with decreasing latitude, but others show the opposite trend (Anstett et al., 2014; Baskett & Schemske, 2018; Kent et al., 2020; Moles et al., 2011; Zvereva & Kozlov, 2021). Various factors such as the community of herbivores, defence trade-offs, and environmental variables have been put forward to explain these contradictory results (Anstett et al., 2014, 2016; Kooyers et al., 2017). We found that standing rates of herbivory increased significantly with latitude. The abundance of ants and larger EFNs that produce more nectar may have led to a strong reduction in herbivory by ant bodyguards despite high herbivore pressure in southern sites. However, we did not find a direct link between ant abundance and herbivory in our SEM, perhaps because of the different time scales reflected in their measurements. The contrasting trends in herbivory and mutualism across latitude in our study suggest that indirect defence mediated by ants or other arthropods may also be an important factor affecting latitudinal clines in herbivory.

This pattern of increasing herbivory with latitude could also have been caused by differences in herbivore abundance and composition across sites rather than patterns in defence investment. Further, plants in northern sites may have been more likely to have escaped from cultivated populations, since Rajasthan and Gujarat, both North Indian states are two of the largest castor-cultivating states in India (Indian Council of Agricultural Research report). Domesticated varieties often show increased susceptibility to herbivory due to defences weakened by generations of artificial selection in managed habitats (Chen et al., 2015; Whitehead et al., 2017 but see Turcotte et al., 2017). This may partially explain the trend of increasing rates of standing herbivory with latitude. This pattern may have also been caused by variation in leaf lifespan across latitude; if northern plants retain leaves for longer, this may lead to higher observed rates of standing herbivory at higher latitudes.

Our results suggest that even a generalized mutualism such as that between ants and EFN-bearing plants can give rise to large-scale latitudinal patterns in species interactions. While a single interaction between an EFN and an ant may not significantly influence the host plant’s fitness, continuous exposure to differing communities of mutualists and antagonists over generations may drive latitudinal patterns. Indeed, our SEM results showed that a snapshot measure of plant fitness, the number of fruits on each plant, decreased with increasing latitude. While environmental factors varying across latitude may have contributed to this effect, mutualism may have played an indirect role in determining host fitness, although we did not see a causal effect of ant abundance or herbivory on the number of fruits in the SEM. Ant-exclusion experiments conducted at different latitudes are needed to understand the contribution of mutualism and herbivory to the fitness of *R. communis*. While snapshot measures are useful in understanding large-scale patterns, long-term studies are crucial, since *R. communis* is a perennial species

Genomic analysis suggests that *R. communis* likely arrived in India around 2000 years ago in an ancient introduction (Xu et al., 2021), thus any adaptation to local mutualist and herbivore fauna occurred is relatively recent. Our data suggest that latitudinal patterns in generalized mutualism may appear over relatively short timescales. The relative contributions of genetic versus environmental variation in mutualism traits across plant populations spanning the latitudinal gradient is not clear from our study. *Ricinus communis* is known to show high phenotypic plasticity in other traits and its success as an introduced species worldwide has been attributed to its ability to tolerate and respond to a wide range of environmental conditions (Goyal et al., 2014; Martins et al., 2011). Likely, EFN traits are also highly plastic, allowing the plant to respond rapidly to local variation in herbivory and mutualist availability. Further, it is known that castor EFN volume is induced by herbivore damage (Wäckers et al., 2001), but much less is known about how other traits like EFN counts and areas respond to herbivory. Common garden studies with controlled breeding designs, and controlled herbivore exposure will help elucidate the source of this variation.

The Indian subcontinent, like much of the Old World tropics, is underrepresented in studies of biodiversity and species interactions (Hargreaves et al., 2020; Millard et al., 2020; Parker et al., 2024; Wan et al., 2020). The Indian subcontinent consists of many diverse ecoregions (Olson & Dinerstein, 1998; Roy et al., 2006), and our study contributes a unique dataset on species interactions of *R. communis* to the literature. Further, our study is one of the first to investigate how an ant-plant mutualism in the same host plant species varies across latitude in both ant and plant partners. While *R. communis* is well-studied in the context of its use as an oilseed, our study highlights the rich natural history of agriculturally important species and their suitability as study systems to address big questions in ecology and evolution. Many agriculturally important species are known to interact with ants and other bodyguards through EFNs, and studies on the ecology of these interactions may help leverage them in the development of sustainable agricultural practices (Jones et al., 2017).

## Open Research Statement

All data and code used in this manuscript are openly available at https://github.com/The-Frederickson-Lab/castor_latitudinal_gradient

## Acknowledgements

We thank Yash Kumar Singhal for his help with measuring EFN areas and volumes, Renee Borges for suggesting *Ricinus communis* as a study system, and Nathan KC for his help with data collection in Port Blair. We would also like to thank Anjani Kammili and Rohit Sasidharan for initial discussions about the study system. For funding, we thank the Natural Sciences and Engineering Research Council of Canada (NSERC) for support via a Discovery Grant to M.E.F.

## Author contributions

PN and MEF conceptualised the study. PN, AS and AD collected the data, and PN performed the analyses and wrote the first draft of the manuscript. AS and AD contributed to data processing.VG provided resources and assistance with fieldwork. All authors reviewed and edited the final draft of the manuscript.

