## Supplementary tables and figures for "Are ant-plant mutualisms stronger at lower latitudes? A case study using the castor plant in the Indian subcontinent"

**Table S1**: Number of plants sampled at each site

| **Site** | **No. of plants sampled** |
| --- | --- |
| Aligarh | 11 |
| Bengaluru | 20 |
| Bhopal | 15 |
| Gwalior | 14 |
| Hampi | 18 |
| Hyderabad | 14 |
| Kanyakumari | 15 |
| Koradi | 14 |
| Port Blair | 10 |
| Ramnagar | 21 |
| Srirangam | 12 |

**Table S2**: Ant species found at each site

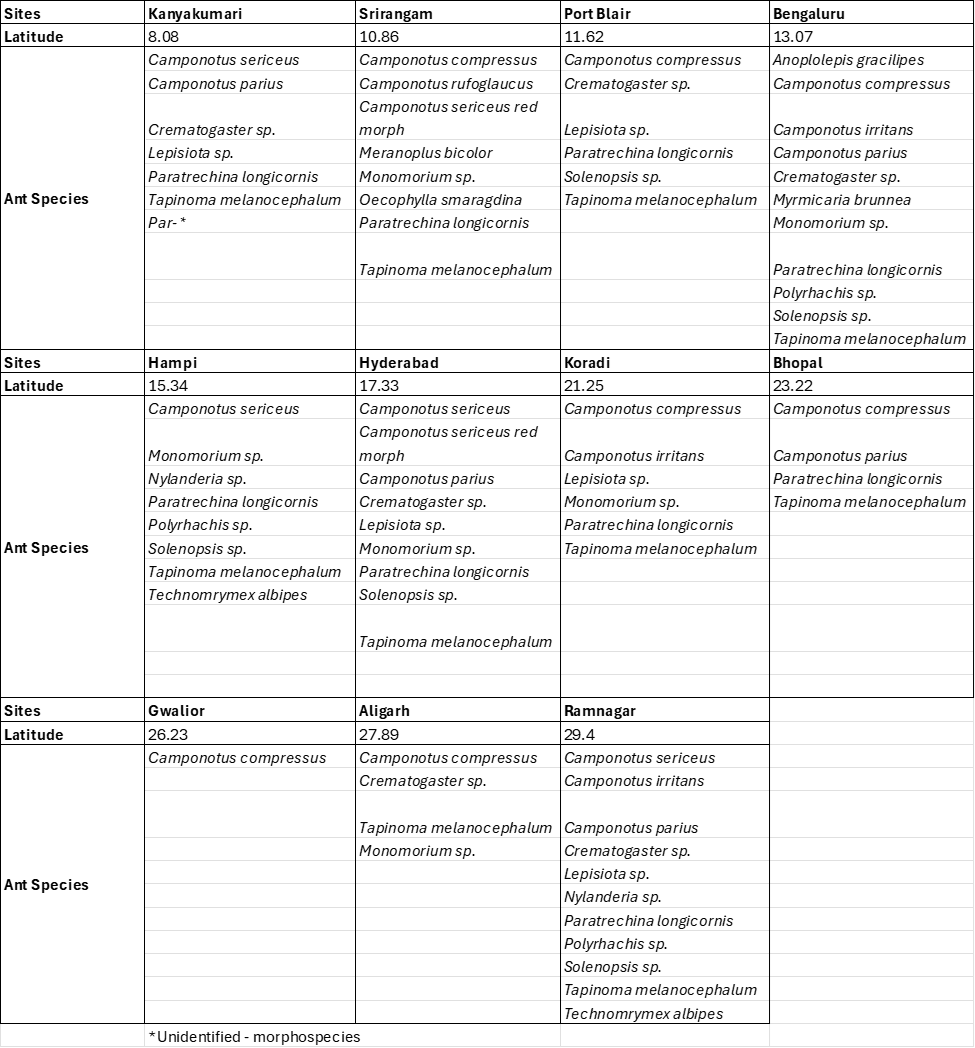

**Table S3**: Linear model results for community characteristics: species richness and Shannon Diversity. Results for the models including all sites (top panel) and all sites except Port Blair, the island site (bottom panel) are shown. Significant effects are italicized and indicated by *** for p ≤ 0:001, ** for p ≤ 0:01 and * for p ≤ 0:05.

| **All sites** | | | | |
| --- | --- | --- | --- | --- |
| Species richness | | | | |
|  | **Estimates** | **SE** | **t value** | **p-value** |
| ***Intercept*** | *8.8126* | *2.8454* | *3.097* | *0.0128 ** |
| **Latitude** | -0.1178 | 0.144 | -0.818 | 0.4344 |
| Shannon diversity | | | | |
|  | **Estimates** | **SE** | **t value** | **p-value** |
| ***Intercept*** | *1.78458* | *0.43088* | *4.142* | *0.00252 *** |
| **Latitude** | -0.02417 | 0.0218 | -1.109 | 0.29633 |
| **Island site excluded** | | | | |
| Species richness | | | | |
|  | **Estimates** | **SE** | **t value** | **p-value** |
| ***Intercept*** | *9.7761* | *3.1551* | *3.099* | *0.0147 ** |
| **Latitude** | -0.1553 | 0.1547 | -1.004 | 0.3447 |
| Shannon diversity | | | | |
|  | **Estimates** | **SE** | **t value** | **p-value** |
| ***Intercept*** | *1.81199* | *0.49528* | *3.658* | *0.00642 *** |
| **Latitude** | -0.02524 | 0.02428 | -1.039 | 0.32897 |

**Table S4**: SEM model results. The standardized estimate is shown for the best fitting model for all variables and covariances. Significant effects are italicized and indicated by *** for p ≤ 0:001, ** for p ≤ 0:01 and * for p ≤ 0:05.

| **Ant visits** | | | | |
| --- | --- | --- | --- | --- |
|  | **Std. estimate** | **SE** | **z value** | **p-value** |
| ***Latitude*** | *-0.377* | *0.078* | *-4.946* | *0**** |
| **Leaf base EFNs** | -0.119 | 0.092 | -1.332 | 0.183 |
| ***Petiole EFNs*** | *0.283* | *0.09* | *3.168* | *0.002*** |
| **Herbivory** | | | | |
|  | **Std. estimate** | **SE** | **z value** | **p-value** |
| ***Latitude*** | *0.312* | *0.082* | *3.759* | *0**** |
| **Ant visits** | -0.108 | 0.078 | -1.33 | 0.183 |
| **Leaf base EFNs** | 0.058 | 0.09 | 0.642 | 0.521 |
| **Petiole EFNs** | 0.088 | 0.386 | 0.946 | 0.344 |
| **Leaf base EFNs** | | | | |
|  | **Std. estimate** | **SE** | **z value** | **p-value** |
| ***Latitude*** | *-0.158* | *0.079* | *-1.998* | *0.046** |
| **Plant age** | -0.136 | 0.079 | -1.723 | 0.085 |
| **Petiole EFNs** | | | | |
|  | **Std. estimate** | **SE** | **z value** | **p-value** |
| **Latitude** | 0.107 | 0.08 | 1.355 | 0.175 |
| **Plant age** | -0.148 | 0.08 | -1.867 | 0.062 |
| **Number of fruits** | | | | |
|  | **Std. estimate** | **SE** | **z value** | **p-value** |
| **Herbivory** | -0.004 | 0.071 | -0.048 | 0.961 |
| **Ant visits** | 0.058 | 0.07 | 0.777 | 0.437 |
| ***Latitude*** | *-0.264* | *0.076* | *-3.319* | *0.001*** |
| **Leaf base EFNs** | -0.068 | 0.08 | -0.821 | 0.412 |
| **Petiole EFNs** | -0.096 | 0.081 | -1.128 | 0.259 |
| ***Plant age*** | *0.376* | *0.066* | *5.442* | *0**** |
| **Covariance** |  |  |  |  |
|  | **Std. estimate** | **SE** | **z value** | **p-value** |
| ***Leaf base ~~ petiole EFNs*** | *0.553* | *0.089* | *6.046* | *0**** |

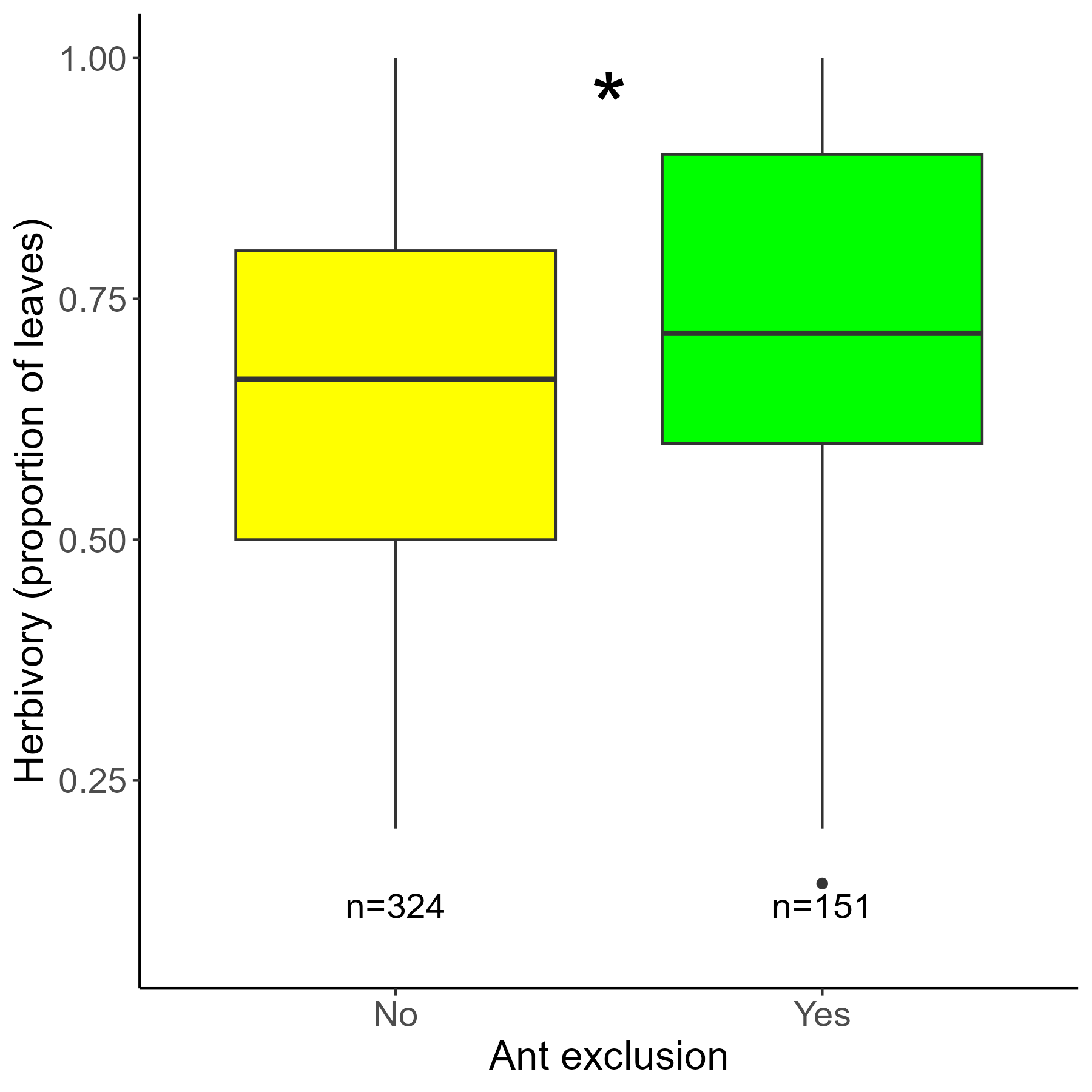

**Figure S1:** The effect of ant exclusion on herbivory in *Ricinus communis*. Results are from an unpublished common garden study using *R. communis* seeds collected from the latitudinal sampling described in this study, in addition to seeds from agricultural varieties (Nathan et al., in prep) in the southern Indian city of Bengaluru (see Fig. 1a). Ant exclusion was implemented by applying a thick layer of Tanglefoot adhesive at the base of each plant’s stem. Plants whose leaves touched the ground or other plants were excluded from this analysis. The asterisk represents a statistically significant effect of the ant exclusion treatment on herbivory in a linear model (p = 6.64e-06). Sample sizes are shown under the boxes.

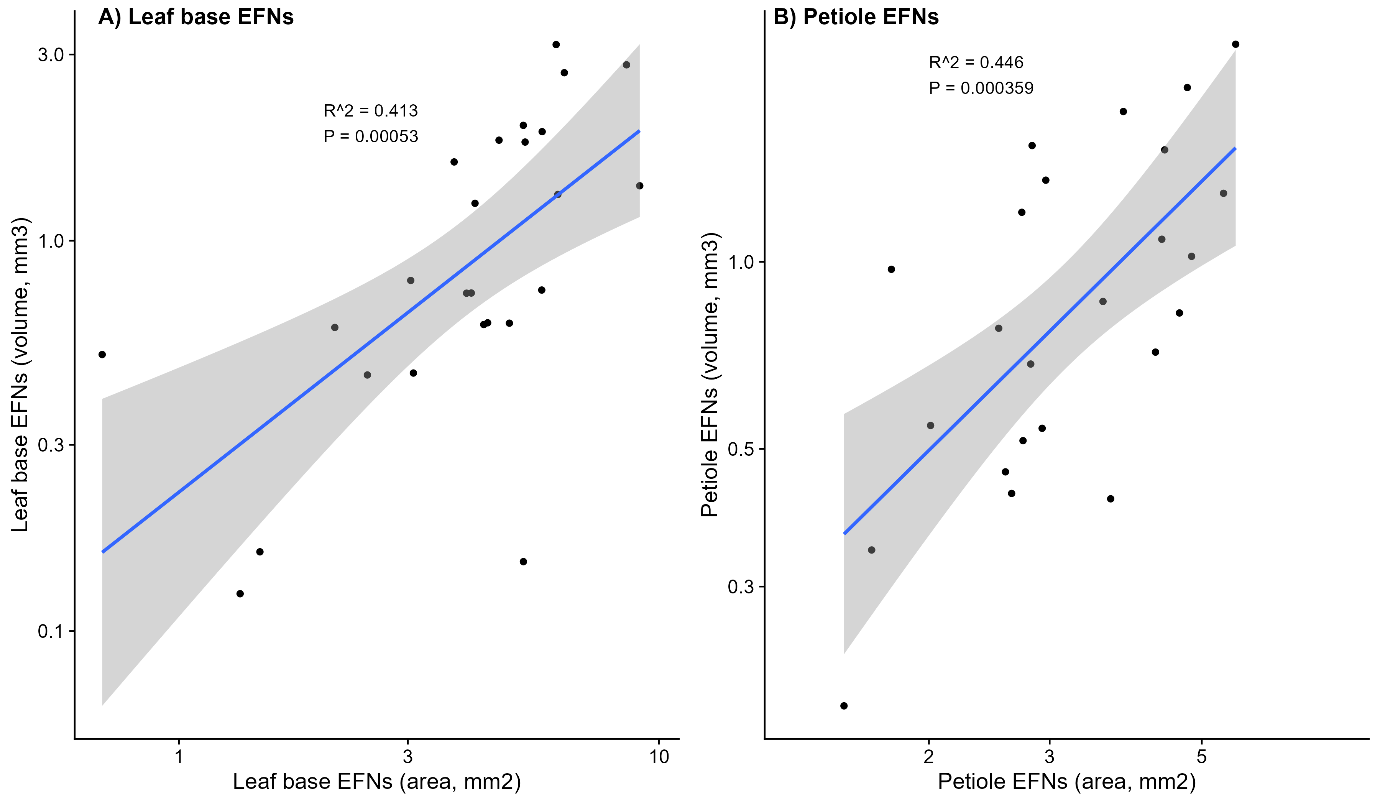

**Figure S2:**  Correlation between EFN areas and volumes. A) Correlation between leaf base EFN areas and volumes. B) Correlation between petiole EFN areas and volumes.

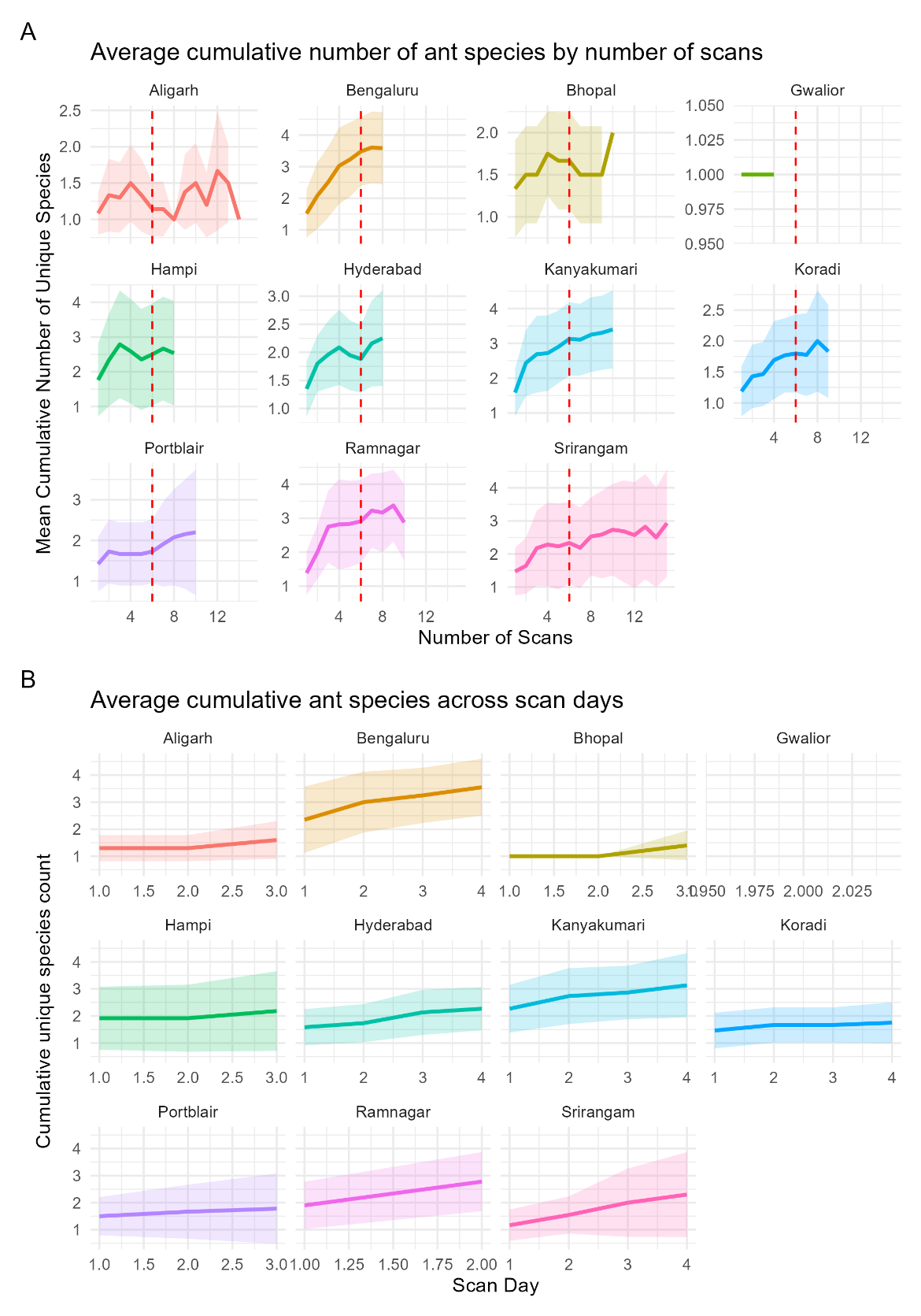

**Figure S3:** Cumulative number of ant species observed across scans and scan days, summarized for each site. A) Average cumulative number of ant species observed across scans, summarized across scan days. The mean number of species across plants is shown as a solid line, with shaded ribbons representing standard deviations. Most sites demonstrate saturation in the number of species observed after approximately 6 scans, as highlighted by the red dashed line. B) Average cumulative number of ant species observed across scan days. The mean cumulative species count summarized across scans for plants in each site is shown as a solid line, with shaded ribbons representing standard deviations. The cumulative number of species increases with additional scan days, indicating that longer observation periods yield more species observations.
